# Natural variation in fertility is correlated with species-wide levels of divergence in *Caenorhabditis elegans*

**DOI:** 10.1101/2021.02.18.431866

**Authors:** Gaotian Zhang, Jake D. Mostad, Erik C. Andersen

## Abstract

Life history traits underlie the fitness of organisms and are under strong natural selection in the face of environmental challenges. A new mutation that positively impacts a life history trait will likely increase in frequency and become fixed in a population (*e.g.* selective sweep). The identification of the beneficial alleles that underlie selective sweeps provides insights into the mechanisms that occurred during the evolution of species. In the global population of *Caenorhabditis elegans,* we previously identified selective sweeps that have drastically reduced chromosomal-scale genetic diversity in the species. Here, we measured the fertility (viable offspring) of a collection of wild *C. elegans* strains, including many recently isolated divergent strains from the Hawaiian islands and found that strains with larger swept genomic regions on multiple chromosomes have significantly higher fertility than strains that do not have evidence of the recent selective sweeps. We used genome-wide association (GWA) mapping to identify three quantitative trait loci (QTL) underlying the fertility variation. Additionally, we mapped previous fertility data of wild *C. elegans* strains and *C. elegans* recombinant inbred advanced intercross lines (RIAILs) that were grown in various conditions and detected eight QTL across the genome using GWA and linkage mappings. These QTL show the genetic complexity of life history traits such as fertility across this species. Moreover, the haplotype structure in each GWA QTL region revealed correlations with recent selective sweeps in the *C. elegans* population. North American and European strains had significantly higher fertility than most strains from Hawaii, a hypothesized origin of the *C. elegans* species, suggesting that beneficial alleles that cause increased fertility could underlie the selective sweeps during the worldwide expansion of *C. elegans.*

## INTRODUCTION

Life history traits are phenotypic characters that affect the fitness of organisms (Knight and Robertson 1957; Stearns 1976, 1989; Charlesworth *et al.* 2003; Flatt and Heyland 2011; Flatt 2020). Traits, such as fertility, size at birth, age at reproductive maturity, and stage- or size-specific rates of survival, interact with each other to affect the fitness of organisms in an ever-changing environment. Genes that affect life history traits should be subject to strong natural selection because they directly affect the fitness of organisms. Adaptive alleles with strong selective advantages in life history related genes are likely to spread rapidly across a population in a selective sweep (Smith and Haigh 1974; Kaplan *et al.* 1989; Berry *et al.* 1991; Stephan 2019). Signatures of selective sweeps include a loss of neutral polymorphism, drastic changes in the site frequency spectrum, and particular patterns of linkage disequilibrium (LD) across the site of selection (Smith and Haigh 1974; Braverman *et al.* 1995; Fay and Wu 2000; Kim and Nielsen 2004; Stephan *et al.* 2006; Stephan 2019). Identification of selective sweeps by these signatures provides a key to locate genes under selection and helps to understand the process of adaptation and evolution.

*Caenorhabditis elegans* is a free-living nematode and a keystone model organism for biological research. The reproductive mode of *C. elegans* is androdioecy, with predominant self-fertilization of hermaphrodites and rare outcrossing between hermaphrodites and males (Brenner 1974). A single hermaphrodite of the laboratory reference strain N2 lays approximately 300 self-fertilized embryos in standard laboratory conditions (Hodgkin and Doniach 1997; Félix and Braendle 2010). Newly hatched animals develop through four larval stages (L1 to L4) into mature reproductive adults after three days in favorable conditions at 20°C (Frézal and Félix 2015). Under stressful conditions, such as crowding and limited food, *C. elegans* enters the dauer diapause stage during larval development to enable survival in harsh environments and to facilitate dispersal. *C. elegans* likely has a boom-and-bust life cycle in the wild because of fluctuating environmental conditions and the spatio-temporal distributed habitats, such as rotting fruits and stems (Félix and Duveau 2012; Frézal and Félix 2015). *C. elegans* is globally distributed (Kiontke *et al.* 2011; Andersen *et al.* 2012; Félix and Duveau 2012; Cook *et al.* 2017; Crombie *et al.* 2019; Lee *et al.* 2020). Although recent studies characterized high genetic diversity of the species in Hawaii and the surrounding Pacific regions (Crombie *et al.* 2019; Lee *et al.* 2020), *C. elegans* exhibits low overall genetic diversity at the global scale (Barrière and Félix 2005; Cutter 2006; Andersen *et al.* 2012). The metapopulation dynamics, seasonal bottlenecks, predominant selfing, low outcrossing rate, low recombination rate, background selection, and recent selective sweeps might all contribute to the low genetic diversity of the species (Barrière and Félix 2005, 2007; Cutter 2006; Rockman and Kruglyak 2009; Rockman *et al.* 2010; Andersen *et al.* 2012). In the genomes of many *C. elegans* strains sampled in temperate regions, chromosomes I, IV, V, and X exhibit signatures of selective sweeps, such as an excess of rare variants, high linkage disequilibrium (LD), and extended haplotype homozygosity over large genomic regions (Andersen *et al.* 2012). By contrast, the genomes of most Hawaiian *C. elegans* strains have no such signatures (Andersen *et al.* 2012; Crombie *et al.* 2019; Lee *et al.* 2020). Analyses of *C. elegans* genetic diversity, population structure, gene flow, and haplotype structure suggest that *C. elegans* originated from the Pacific region, such as the Hawaii Islands, the western United States, or New Zealand, and expanded worldwide, especially into human-associated habitats (Andersen *et al.* 2012; Crombie *et al.* 2019; Lee *et al.* 2020). The recent positive selective sweeps likely occurred during this expansion, but the beneficial alleles that have driven the sweeps and their fitness advantages are yet unknown.

Here, we measured lifetime fertility of 121 wild *C. elegans* strains and compared this trait between swept strains that experienced the recent selective sweeps and divergent strains that avoided these sweeps. We found that swept strains had significantly higher lifetime fertility than divergent strains, as well as significant geographical differences in lifetime fertility between strains from the Hawaii Islands and strains from other parts of the world. We then used GWA mapping to identify three QTL on chromosome I, II, and V that influence the lifetime fertility of *C. elegans.* Additionally, we identified eight QTL impacting *C. elegans* fertility in different environments using GWA and linkage mappings of previous fertility data. The 11 QTL reveal the complex genetic architecture of *C. elegans* fertility. Furthermore, we discovered that the different alleles at each QTL peak marker and the different haplotypes in each QTL among the 121 strains were strongly correlated with signatures of recent selective sweeps found in each strain. Our results suggest that higher lifetime fertility could have provided selective advantages for swept strains and the underlying genetic variants might have driven the recent strong sweeps in the *C. elegans* strains that have colonized the world.

## MATERIALS AND METHODS

### *C. elegans* strains

All the wild strains were obtained from *C. elegans* Natural Diversity Resource (CeNDR) (Cook *et al.* 2017). Animals were cultured at 20°C on modified nematode growth medium (NGMA) containing 1% agar and 0.7% agarose to prevent burrowing and fed *Escherichia coli (E. coli)* strain OP50. Prior to each assay, strains were grown for three generations without entering starvation or encountering dauer-inducing conditions (Andersen *et al.* 2014).

### Swep2b haplotypes and strains

Haplotype data for 403 *C. elegans* isotypes, representing 913 wild strains, were acquired from the 20200815 CeNDR release. We compared the total length of each haplotype per chromosome across all isotypes to identify the most common haplotypes on each chromosome. We then searched for the regions of the most common haplotypes in each *C. elegans* isotype and recorded them if their length was greater than 1 Mb (Crombie *et al.* 2019; Lee *et al.* 2020). We classified haplotypes outside of recorded regions as unswept haplotypes. The swept status of some haplotypes was undetermined when no identical-by-descent groups were found, and thus the haplotype information for that region was missing in the CeNDR release.

Signatures of selective sweeps were identified on chromosomes I, IV, V, and X, but not on chromosomes II and III (Andersen *et al.* 2012). Therefore, we focused on the four chromosomes (I, IV, V, and X) and defined their most common haplotypes as swept haplotypes (Lee *et al.* 2020). In each *C. elegans* isotype, chromosomes that contain greater than or equal to 30% of the swept haplotype were classified as swept chromosomes. We classified isotypes with any swept I, IV, V, and X chromosomes as swept isotypes and isotypes without any swept I, IV, V, and X chromosomes as divergent isotypes. Strains that belong to swept isotypes and divergent isotypes were classified as swept strains and divergent strains, respectively (Gilbert *et al.* 2020).

### Genetic relatedness

Genetic variation data for 403 *C. elegans* isotypes were acquired from the hard-filtered isotype variant call format (VCF) 20200815 CeNDR release. These variants were pruned to the 1,074,596 biallelic single nucleotide variants (SNVs) without missing genotypes. We converted this pruned VCF file to a PHYLIP file using the vcf2phylip.py script (Ortiz 2019). The unrooted neighbor-joining tree was made using the R packages phangorn (v2.5.5) and ggtree (v1.14.6) (Schliep 2011; Yu *et al.* 2017).

### Fertility measurements

For each *C. elegans* strain, single L4 larval stage hermaphrodites were picked to each of five 3.5 cm plates with NGMA and OP50, and were maintained at 20°C. For each assay plate, the original hermaphrodite parent was transferred to a fresh plate every 24 hours for 96 hours. A custom-built imaging platform (DMK 23GP031 camera; Imaging Source, Charlotte, NC) was used to collect images for each of the first four assay plates (0, 24, 48, and 72 hour samples) 48 hours after removal of the parent from each plate. Most strains had few offspring after 96 hours. Images of the fifth assay plates were collected 72 hours after the final transfer of the parents. From each image, the total offspring was counted by visual inspection using the Multi-point Tool in ImageJ (v1.8.0_162) (Schneider *et al.* 2012). The original hermaphrodite parents on the fifth assay plates were excluded from the counts. The number of offspring in each of the first four assay plates corresponds to the daily fertility. Numbers of offspring on the fifth assay plates contained offspring from three days. For each biological replicate of each *C. elegans* strain, the lifetime fertility was calculated as the total number of offspring from the five plates. Few parent animals died during the assays. Only biological replicates with data from all five assay plates were used in the calculations of daily and total fertility. We collected fertility data for 557 replicates of 121 *C. elegans* strains (mean lifetime fertility (MLF) = 231, standard deviations (SD) = 55): 84 strains with five replicates (MLF = 232, SD = 55), 28 strains with four replicates (MLF = 229, SD = 52), seven strains with three replicates (MLF = 214, SD = 49), and two strains with two replicates (MLF = 292, SD = 19).

### Genome-wide association (GWA) mapping

GWA mapping was performed on the mean fertility measurements of biological replicates from 121 *C. elegans* strains, which belong to 121 distinct isotypes. Genotype data for each of the 121 isotypes were acquired from the hard-filtered isotype VCF (20200815 CeNDR release). We performed the mapping using the pipeline cegwas2-nf (https://github.com/AndersenLab/cegwas2-nf) as previously described (Zdraljevic *et al.* 2019; Na *et al.* 2020). Briefly, we used BCFtools (Li 2011) to filter variants that had any missing genotype calls and variants that were below the 5% minor allele frequency. We used PLINK v1.9 (Purcell *et al.* 2007; Chang *et al.* 2015) to prune the genotypes to 56,878 markers with a linkage disequilibrium (LD) threshold of *r^2^* < 0.8 and then generated the kinship matrix using the *A.mat* function in the R package rrBLUP (v4.6.1) (Endelman 2011). The number of independent tests (*N_test_*) within the genotype matrix was estimated using the R package RSpectra (v0.16.0) (https://github.com/yixuan/RSpectra) and correlateR (0.1) (https://github.com/AEBilgrau/correlateR). The eigen-decomposition significance (EIGEN) threshold was calculated as −log_10_(0.05/*N_test_*). We used the *GWAS* function in the rrBLUP package to perform the genome-wide mapping with the EMMA algorithm (Kang *et al.* 2008). QTL were defined by at least one marker that was above the Bonferroni-corrected significance (BF) threshold, to locate the best estimate of QTL positions with the highest significance. We used the *LD* function from the R package genetics (v1.3.8.1.2) (https://cran.r-project.org/package=genetics) to calculate the LD correlation coefficient *r^2^* among the QTL peak markers associated with *C. elegans* lifetime fertility.

We also performed GWA mapping using fertility data in DMSO control conditions from a previous study (Hahnel *et al.* 2018), where 236 *C. elegans* wild strains were cultured and phenotyped using the high-throughput fitness assays (HTA) as previously described. Briefly, L4 larval stage hermaphrodites were cultured to gravid adult stage on plates and were bleached to obtain synchronized offspring. The embryos were grown to L4 larval stage in liquid (K medium) (Boyd *et al.* 2012) and fed an *E. coli* HB101 lysate (García-González *et al.* 2017) in 96-well plates. A large-particle flow cytometer (COPAS BIOSORT; Union Biometrica, Holliston, MA) was used to sort three L4 larvae into each well of new 96-well plates containing K medium, *E. coli* HB101 lysate, and 1% DMSO. Animals in the 96-well plates were incubated at 20°C for 96 hours to allow animals to grow and produce offspring, followed by measurements of various fitness parameters, including fertility. Raw fertility data were pruned, normalized, and regressed using the R package *easysorter* (v1.0) (Shimko and Andersen 2014; Hahnel *et al.* 2018). The processed fertility, norm.n, of each strain was used here for GWA mapping.

### Statistical analysis

Statistical significance of fertility differences between swept strains (groups) and divergent strains (groups), and fertility differences among different sampling locations, was tested with the Wilcoxon test using the *stat_compare_means* function in the R package ggpubr (v0.2.4) (https://github.com/kassambara/ggpubr/). Broad-sense heritability of *C. elegans* lifetime fertility was calculated using the *lmer* function in the R package lme4 (v1.1.21) with the model *phenotype ~ 1+* (*1\strain*) (Bates *et al.* 2015).

### Linkage mapping

We performed linkage mapping using fertility data from a large panel of recombinant inbred advanced intercross lines (RIAILs) derived from QX1430 and CB4856 (Andersen *et al.* 2015). The fertilities (norm.n) of the RIAILs and the parents were measured using the HTA as described above, under three conditions: 1% H2O (402 RIAILs), 1% DMSO (417 RIAILs), and 0.5% DMSO (432 RIAILs). Linkage mapping was performed on each trait using the R package *linkagemapping* (v1.3) (https://github.com/AndersenLab/linkagemapping) and the single-nucleotide variation data of the RIAILs in the package as described previously (Evans and Andersen 2020). Briefly, logarithm of the odds (LOD) scores for each genetic marker and each trait were calculated using the function *fsearch.* The QTL threshold for significant LOD scores in each mapping was defined by permuting trait values 1000 times, mapping the permuted trait data, and taking the 95th quantile LOD score as the 5% genome-wide error rate. 95% confidence intervals of each QTL were determined using the function *annotate_lods.*

### Data availability

File S1 contains the haplotype data of 403 *C. elegans* isotypes from CeNDR release 20200815. File S2 contains genetic relatedness of 403 *C. elegans* isotypes. File S3 contains lifetime fertility of 121 *C. elegans* strains and their classification of swept strains and divergent strains. File S4 contains daily fertility of 121 *C. elegans* strains. File S5 contains GWA results on lifetime fertility of 121 *C. elegans* strains. File S6 contains genotype and phenotype data of 121 *C. elegans* strains at the peak markers of GWA mapping. File S7 contains the sampling locations of 121 *C. elegans* strains. File S8 contains the GPS coordinates of sampling locations of 121 *C. elegans* strains. File S9 contains lifetime fertility and swept and divergent classifications of each of the four swept chromosomes for each of the 121 *C. elegans* strains. File S10 contains LD results among the three QTL of GWA using 121 *C. elegans* strains. File S11 contains the shared haplotypes of the 121 strains within the QTL of GWA mapping. File 12 contains GWA results on fertility data of 236 strains from a previous study (Hahnel *et al.* 2018). File S13 contains genotype and phenotype data of 236 strains at the peak marker of GWA mapping. File S14 contains the shared haplotypes of the 236 strains within the QTL of GWA mapping. File S15 contains the linkage mapping results for the 402 RIAILs in 1% water condition. File S16 contains genotype and phenotype data of the 402 RIAILs at the peak markers and phenotype data of the parents in linkage mapping results. File S17 contains the linkage mapping results for the 417 RIAILs in 1% DMSO condition. File S18 contains genotype and phenotype data of the 417 RIAILs at the peak markers and phenotype data of the parents in linkage mapping results. File S19 contains the linkage mapping results for the 432 RIAILs in 0.5% DMSO condition. File S20 contains genotype and phenotype data of the 432 RIAILs at the peak markers and phenotype data of the parents in linkage mapping results. The datasets and code for generating figures can be found at https://github.com/AndersenLab/swept_broods.

## RESULTS

### Chromosome-scale sweeps shape *C. elegans* strain relationships

Genomic information of 913 wild *C. elegans* strains, grouped into 403 genetically distinct isotypes, are currently available in *C. elegans* Natural Diversity Resource (CeNDR) (Cook *et al.* 2017). The latest CeNDR haplotype data, inferred from identical-by-descent groups among the 403 isotypes, include 22,859 distinct haplotypes across the genome. The number of haplotypes on each chromosome ranged from 2,567 to 5,199. We identified 11 most common haplotypes found in the majority of wild strains. Of the 403 *C. elegans* isotypes, 331 share more than 1 Mb of regions with at least one of the 11 most common haplotypes, particularly on chromosomes I, IV, V and X (Figure 1A, File S1). The haplotype structure of shared haplotypes over large regions across 403 isotypes further supported the selective sweeps identified previously (Andersen *et al.* 2012).

**Figure 1.**
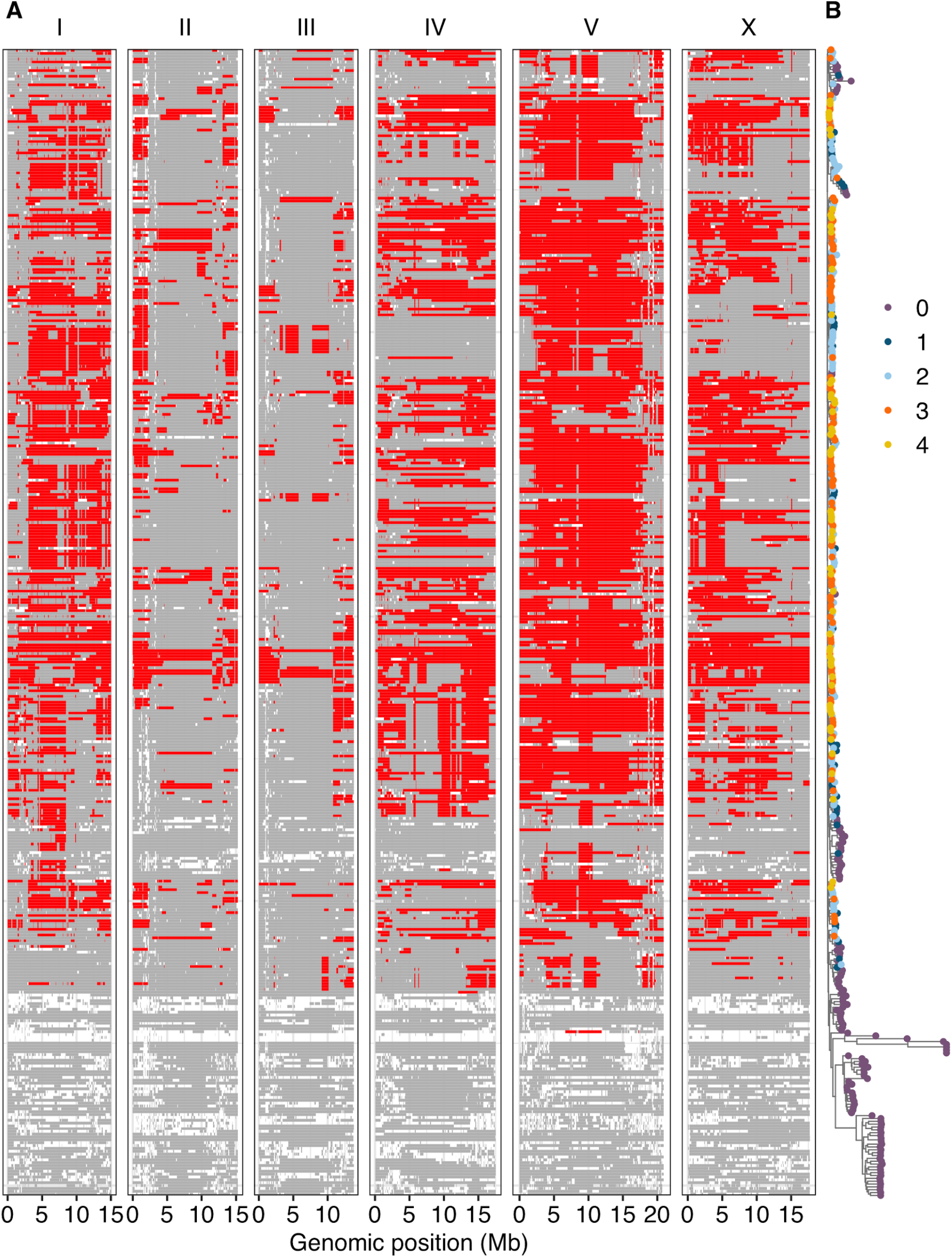
Swept chromosomes and genetic relatedness of wild *C. elegans* isotypes. (A) Sharing of the most common haplotypes (red) across the genome of *C. elegans* for 403 isotypes is shown. Genomic regions of unswept haplotypes (haplotypes other than the most common haplotypes) are colored gray. White segments are undetermined haplotypes in regions where no identical-by-descent groups were found (Crombie *et al.* 2019). The genomic position is plotted on the x-axis. Each row on the y-axis represents one of the 403 isotypes, ordered as their positions in (B). (B) A tree showing genetic relatedness of the 403 *C. elegans* isotypes, using 1,074,596 biallelic segregating sites, is shown. The tips of the tree are colored by the number of swept chromosomes (purple for zero, deep blue for one, light blue for two, orange for three, and gold for four) in each *C. elegans* isotype.

The shared fraction of the most common haplotypes per chromosome varies in each *C. elegans* isotype. Among chromosomes with shared regions in the 331 isotypes, chromosomes I, II, III, IV, V and X have mean shared fractions and SD of 0.45 ± 0.25, 0.21 ± 0.19, 0.22 ± 0.17, 0.52 ± 0.28, 0.60 ± 0.27, and 0.43 ± 0.28, respectively. We focused on swept haplotypes, the most common haplotypes on chromosomes I, IV, V and X, where evidence of selective sweeps were identified (Andersen *et al.* 2012). The chromosomal sharing of swept haplotypes contributes substantially to the genetic relatedness of *C. elegans* isotypes (Figure 1B, File S2). Isotypes with swept chromosomes, which contain greater than or equal to 30% of swept haplotypes, clustered together. Of the 331 isotypes noted above, 281 have at least one swept chromosome (Figure 1B). We classified these 281 *C. elegans* isotypes as swept isotypes. We found that 244 swept isotypes have at least two swept chromosomes. By contrast, most of the 122 divergent isotypes with no swept chromosomes clustered together (Figure 1B). Previous analyses on genome-wide average nucleotide diversity (π), Tajima’s *D,* and genome-wide Hudson’s *Fst* between 43 Hawaiian isotypes (most are divergent isotypes) and 233 non-Hawaiian isotypes (most are swept isotypes) also revealed a high degree of divergence, the highest of which were found in genomic regions impacted by the selective sweeps (Crombie *et al.* 2019). The high degree of genetic relatedness across the species is driven by the selective sweeps, but the fitness advantage causing the strong selective sweeps is yet unknown.

### Natural variation in fertility among swept and divergent strains

To compare the fitness between swept and divergent isotypes, we measured lifetime fertility of 121 wild *C. elegans* strains sampled across the globe (Figure S1, File S8). Single fourth larval stage hermaphrodites were transferred daily for five days and maintained under normal laboratory conditions. We manually counted the viable offspring from images of assay plates. The results showed large variation in lifetime fertility among wild *C. elegans* strains (Figure 2A, File S3). The mean lifetime fertility ranged from 106 to 335 offspring among the 121 strains. We observed the species reproductive peak in the second day of the assay, with a median peak number of 109 offspring (Figure 2B, File S4).

**Figure 2.**
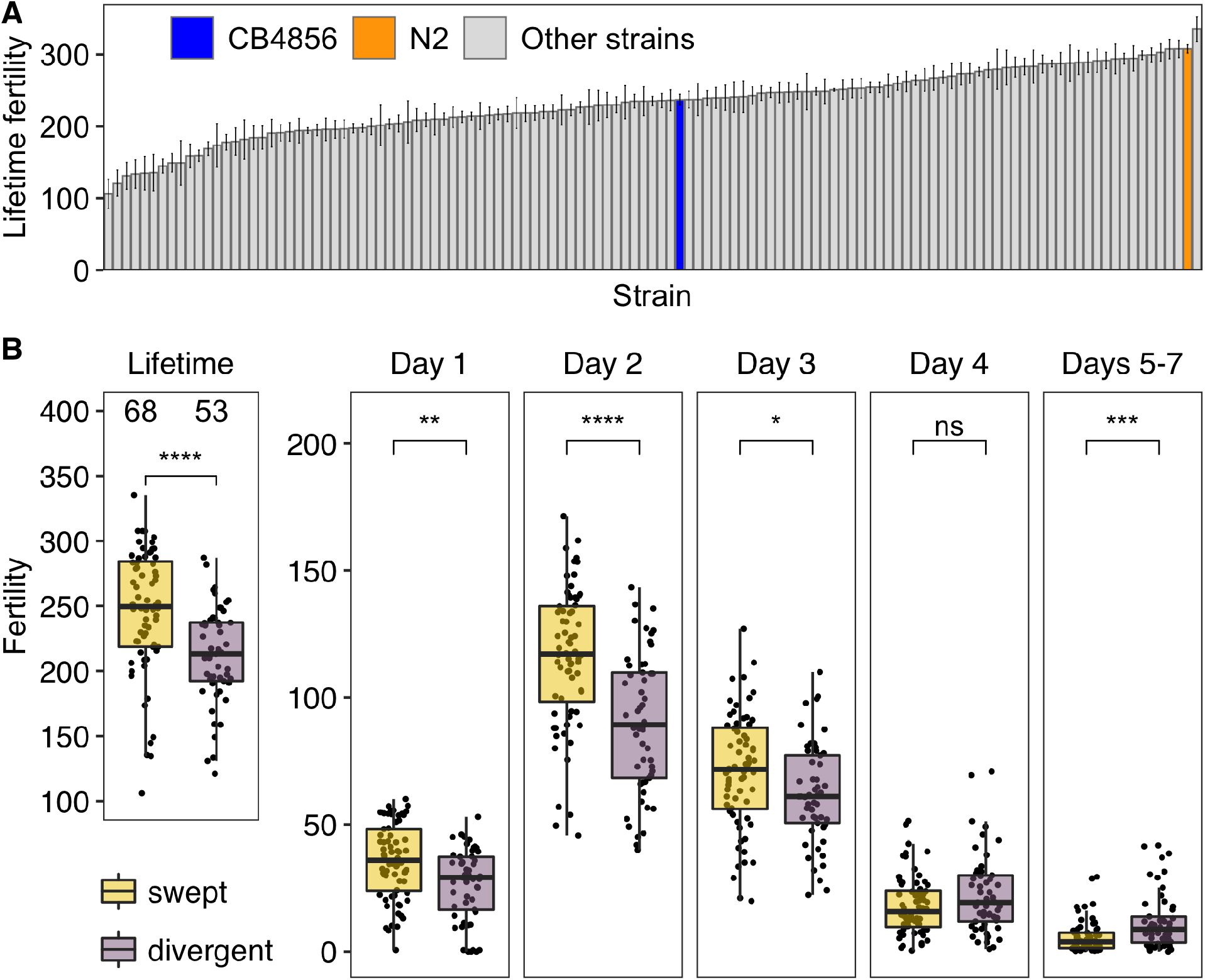
Natural variation in *C. elegans* fertility. (A) A bar plot for lifetime fertility (y-axis) of 121 wild *C. elegans* strains is shown. Strains on the x-axis are sorted by their mean lifetime fertility of two to five biological replicates. Error bars show standard errors of lifetime fertility among replicates. The lab reference strain N2 and the Hawaii strain CB4856 are colored orange and blue, respectively; other strains are colored gray. (B) Comparisons of lifetime and daily fertility between 68 swept strains (gold) and 53 divergent strains (purple) are shown as Tukey box plots. Statistical significance was calculated using the Wilcoxon test. Significance of each comparison is shown above each comparison pair (ns: p-value > 0.05; *: p-value ≤ 0.05; **: p-value ≤ 0.01; ***: p-value ≤ 0.001; ****: p-value ≤ 0.0001).

Of the 121 *C. elegans* strains, 68 strains were classified as “swept” strains and 53 strains were classified as “divergent” strains (see Methods, Figure 2B, Figure S1, File S3). Mean lifetime fertility of swept strains was significantly higher than divergent strains (Wilcoxon test, *p* = 9.1E-6) (Figure 2B). Because different strains could have different swept chromosomes, we extended the comparisons to chromosome levels (Figure S2, File S9). We assigned strains into swept groups or divergent groups in each swept chromosome, depending on whether isotypes had a specific swept chromosome.

Although the numbers of strains in the two groups were different across swept chromosomes, swept groups always showed significantly higher lifetime fertility than divergent groups (Wilcoxon test, *p* < 0.0001) (Figure S2). The striking differences in lifetime fertility suggested that swept strains have higher fitness than divergent strains under normal laboratory conditions. Additionally, we compared the daily fertility between swept and divergent strains. We found that swept strains showed significantly higher daily fertility than divergent strains in the first three days of the assays (Wilcoxon test, *p* = 0.0016, *p* = 1.7E-6, and *p* = 0.014, respectively) (Figure 2B). This significant difference of fertility between swept and non-swept groups provided an opportunity to dissect the genetic basis of the natural variation in lifetime fertility. We calculated the broad-sense heritability and found a substantial heritable genetic component (*H^2^* = 0.63) of the phenotypic variance across these strains.

### Three QTL are associated with natural variation in *C. elegans* lifetime fertility

To identify genomic loci that underlie fertility variation, we performed a marker-based GWA mapping using mean lifetime fertility data from 121 *C. elegans* strains and the whole-genome variant data from CeNDR. We identified three distinct QTL (Figure 3A, File S5). The first QTL, located on the right arm of chromosome I, has a peak-marker at position 13,917,228 and explains 21% of the phenotypic variation among the 121 strains. The second QTL located on the left arm of chromosome II has a peak-marker position at 543,326 and explains 22% of the phenotypic variation. The third QTL spans the center of chromosome V with the peak marker located at 14,534,671 and explains 30% of the phenotypic variation. Because of the strong LD within and between chromosomes in *C. elegans* (Andersen *et al.* 2012), linked regions might be falsely discovered as QTL even though they have no variants that underlie the phenotypic variation. To test the independence of the three QTL, we calculated the pairwise LD among their peak markers (Figure S3, File S10). The results showed moderate levels of LD (ranged from 0.387 to 0.512) for all three pairs, suggesting that they might not be independent. Notably, at all QTL peak markers, most swept strains have the reference alleles and most divergent strains have the alternative alleles (Figure 3B, File S6). We further compared the sharing of haplotypes among the 121 strains within each QTL region (Figure S4, File S11). The majority of the strains with the reference alleles at the peak markers have the most common haplotypes in the QTL regions. By contrast, few strains with alternative alleles have the most common haplotypes in the QTL regions. Taken together, these results suggest that the genetic variants and different haplotypes underlying lifetime fertility variation might be linked to the selective sweeps in the global population of *C. elegans.*

**Figure 3.**
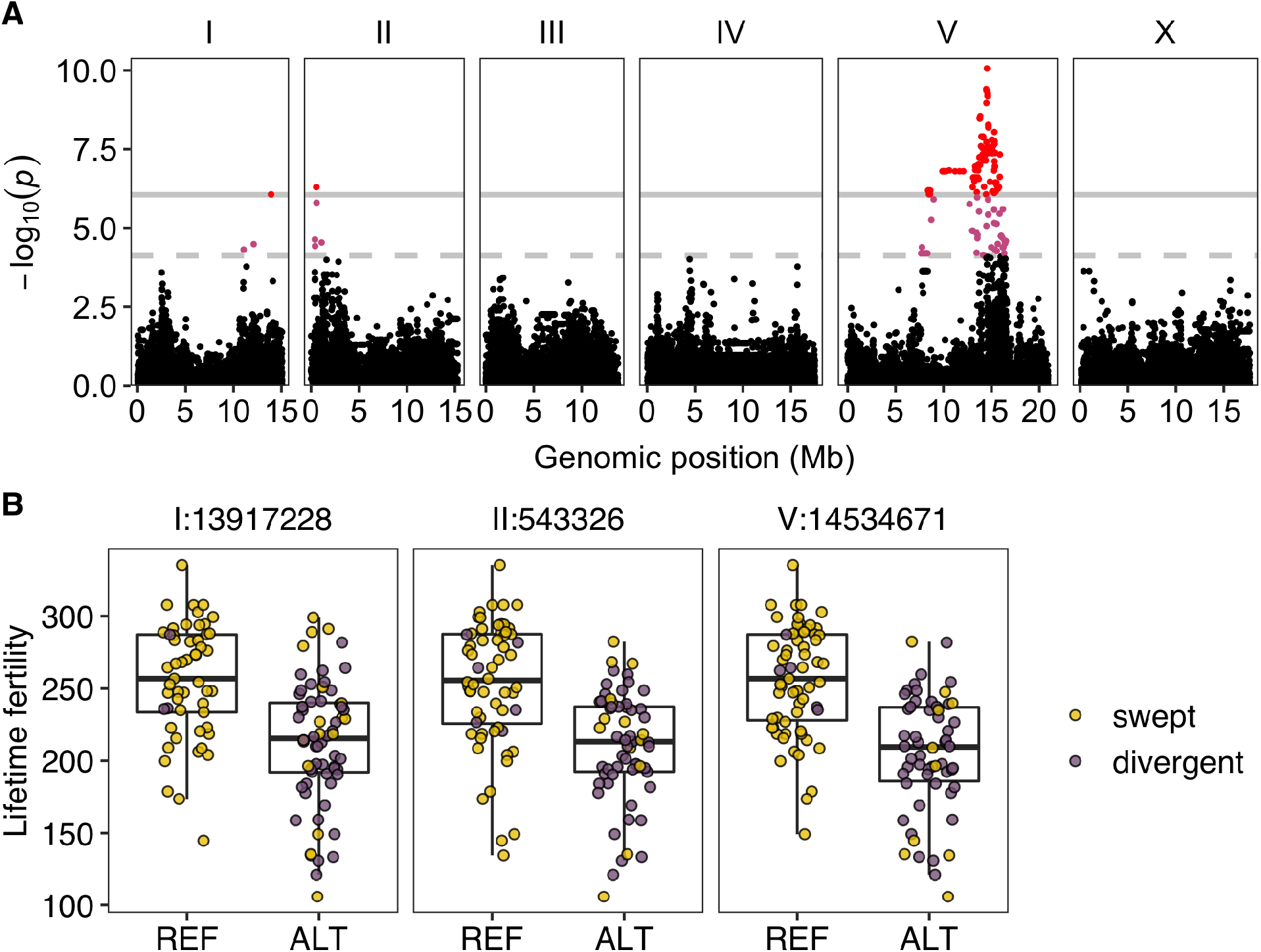
Three QTL were identified in GWA mapping of lifetime fertility variation in 121 *C. elegans* wild strains. (A) Manhattan plot indicating GWA mapping results. Each point represents an SNV that is plotted with its genomic position (x-axis) against its −log_10_(p) value (y-axis) in mapping. SNVs that pass the genome-wide EIGEN threshold (the dotted gray horizontal line) and the genome-wide BF threshold (the solid gray horizontal line) are colored pink and red, respectively. (B) Tukey box plots showing lifetime fertility between strains with different genotypes at the peak marker position in each QTL. Each point corresponds to a *C. elegans* strain and is colored gold for swept strains and purple for divergent strains. On the x-axis, REF represents strains with the N2 reference allele and ALT represents strains with the alternative allele.

### Hawaiian *C. elegans* exhibit lower lifetime fertility than strains sampled across the globe

Most of the 121 *C. elegans* strains were originally sampled from three geographically isolated locations: 50 from the Hawaiian Islands, 22 from North America, and 41 from Europe (Figure S1). Of the 50 Hawaiian *C. elegans* strains, 46 were classified as divergent, and the other four strains have no more than two swept chromosomes (Figure 4, Figure S1, File S7). Most *C. elegans* strains from North America and Europe were classified as swept strains (Figure 4, Figure S1, File S7). We compared lifetime fertilities of strains isolated from these three locations (Figure 4). Compared to strains from North America and Europe, Hawaiian strains had significantly lower lifetime fertility (Wilcoxon test, *p* = 0.00063 and *p* = 7.5E-6, respectively). The difference in lifetime fertility between strains from North America and strains from Europe was insignificant. These data suggested that the selective sweeps that occurred outside Hawaii contribute substantially to the geographical lifetime fertility difference.

**Figure 4.**
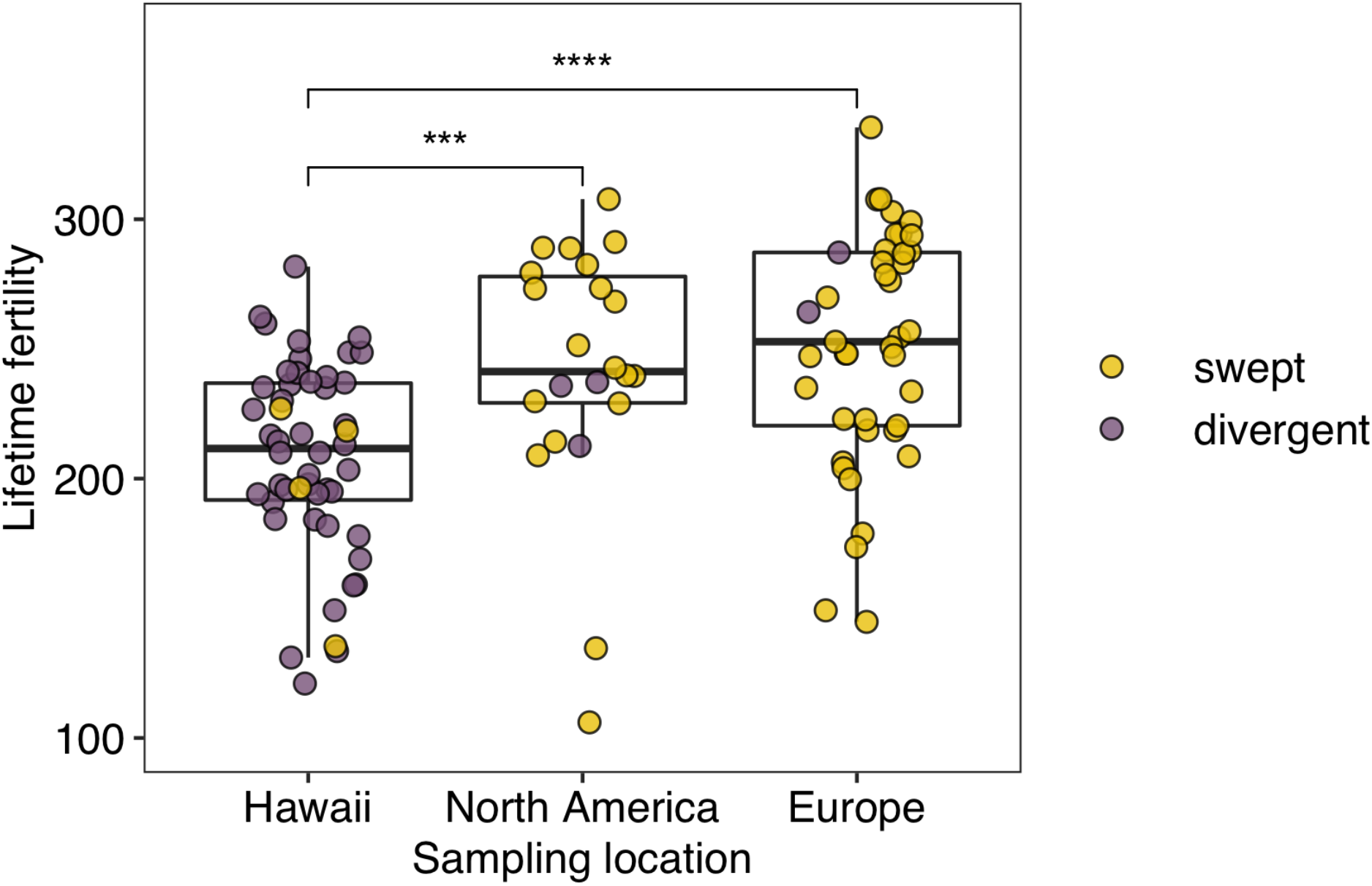
Lifetime fertility comparisons in wild *C. elegans* strains among different sampling locations. Comparisons of lifetime fertility among strains collected from Hawaii (50 strains), North America (22 strains), and Europe (41 strains). Each point corresponds to a strain and is colored gold for swept strains and purple for divergent strains. Statistical significance was calculated using the Wilcoxon test. Significance of each comparison is shown above each comparison pair (***: p-value ≤ 0.001; ****: p-value ≤ 0.0001). The difference of lifetime fertility between North American and European strains is insignificant.

### More QTL underlying lifetime fertility of *C. elegans*

We also mapped the fertility data in the 1% DMSO control condition from one of our published studies that used the high-throughput fitness assays (HTA) (See Methods) to measure various fitness parameters of 236 strains (209 swept strains and 27 divergent strains) (Hahnel *et al.* 2018). Here, we performed GWA mapping using the fertility measurements (norm.n) and identified a QTL on chromosome X (from 3.9 Mb to 5.4 Mb, with the peak marker at 4,831,537) (Figure 5, Figure S5A, File S12). Divergent strains showed no enrichment with either genotype at the peak marker (Figure S5B, File S13). However, most strains with the reference allele have the most common haplotypes and most strains with the alternative allele have unswept haplotypes (Figure S5C, File S14). These results suggest that the genetic variants in this region might also be linked to the recent selective sweeps in wild *C. elegans* populations.

**Figure 5.**
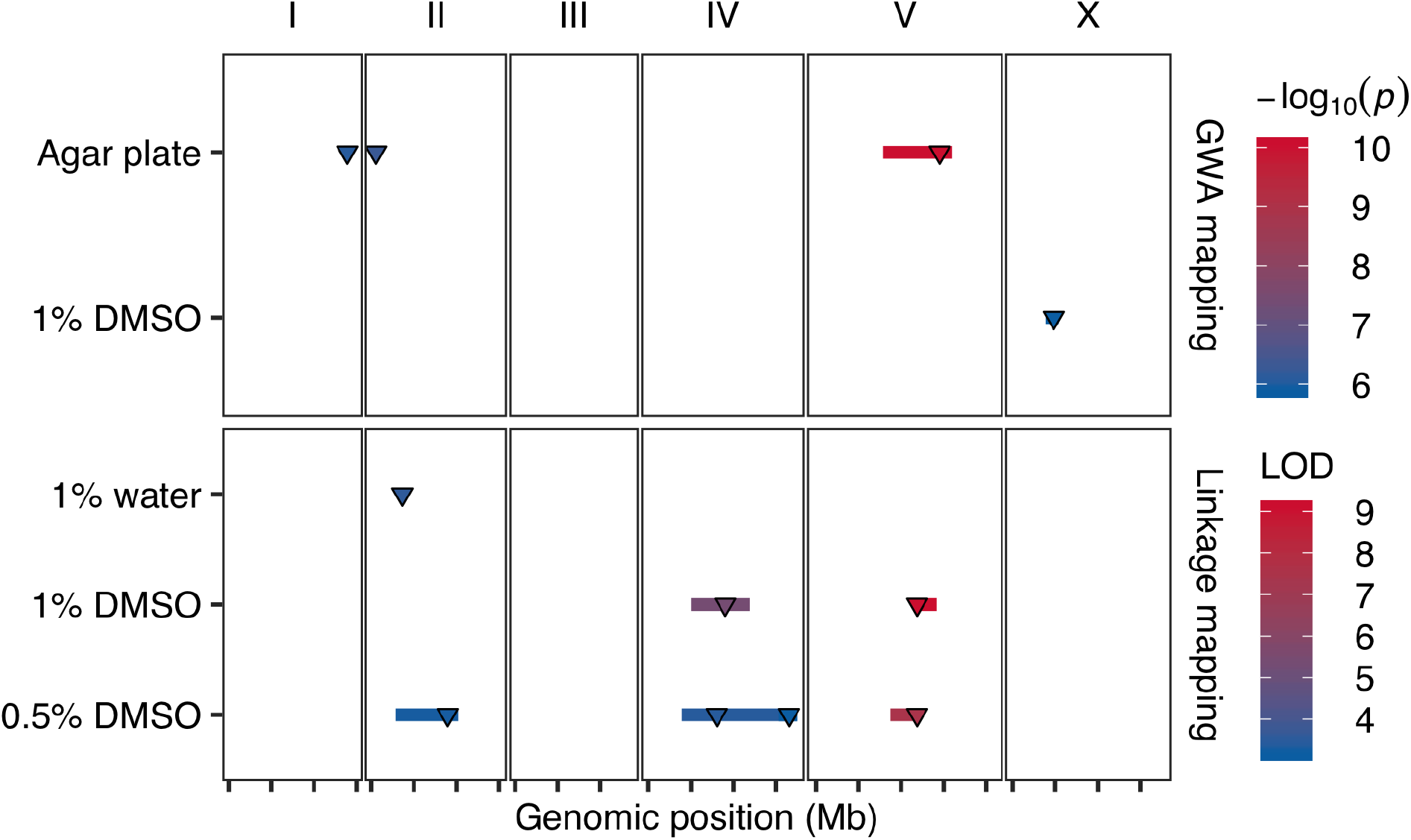
Multiple QTL impacting *C. elegans* lifetime fertility in different conditions. Four GWA mapping QTL of two conditions (121 strains cultured in agar plate and 236 strains cultured in liquid with 1% DMSO) and seven linkage mapping QTL of three conditions (*C. elegans* RIAILs cultured in liquid with 1% water, 1% DMSO, and 0.5% DMSO, respectively) are plotted. Each condition is plotted on the y-axis against the genomic position of its QTL on the x-axis separated by chromosomes with tick marks denoting every 5 Mb. Each QTL is plotted as a line with a triangle indicating the peak marker and colored by the −log_10_(p) value (GWA QTL) or the logarithm of the odds (LOD) score (for linkage mapping QTL), increasing in significance from blue to red.

Also using HTA as above, we measured fertility in liquid culture using the *C. elegans* recombinant inbred advanced intercross lines (RIAILs) derived from QX1430 and CB4856 (Andersen *et al.* 2015) under three conditions: 1% water, 1% DMSO, and 0.5% DMSO (see Methods). In contrast to the fertility variation of *C. elegans* strains cultured in agar plates, the N2 strain showed lower fertility than the CB4856 strain using HTA (Figure S6B, Figure S7B, Figure S8B, File S16, File S18, File S20), indicating that environmental factors can have drastic effects on *C. elegans* fertility. We found seven QTL for fertility on chromosomes II, IV, and V under the three conditions (Figure 5, Figure S6A, Figure S7A, Figure S8A). In 1% water, linkage mapping identified a single QTL confidence interval (II: 3.4 Mb - 4 Mb) on the left arm of chromosome II (Figure 5, Figure S6A, File S15). In 1% DMSO, linkage mapping identified two QTL located on chromosomes IV (5 Mb - 11.9 Mb) and V (11.8 Mb - 14.2 Mb), respectively (Figure 5, Figure S7A, File S17). In 0.5% DMSO, the four QTL on chromosomes II (2.9 Mb - 10.2 Mb), IV (3.9 Mb - 17.5 Mb), and V (8.7 Mb - 12.3 Mb) recapitulated the three QTL detected in 1% water and 1% DMSO, respectively (Figure 5, Figure S8A, File S19). Furthermore, the QTL on chromosome V in both DMSO conditions overlapped with the GWA QTL on chromosome V using the 121 wild strains in agar plates (Figure 5). Because linkage mapping using this set of *C. elegans* RIAILs can only find QTL variants in the CB4856 strain, overlapping of QTL between linkage mapping and GWA mapping suggests that the CB4856 strain carries the common alternative alleles among wild *C. elegans* strains in the shared regions. Altogether, these results suggest that *C. elegans* might have shared and separated loci controlling fertility in agar cultures and in liquid cultures with slightly different concentrations of DMSO.

## DISCUSSION

In this study, we report natural variation of lifetime fertility for 121 wild *C. elegans* strains and found that the previously reported chromosome-scale selective sweeps play a key role in the different fertilities among strains. We defined swept haplotypes, swept isotypes, and swept strains, using the latest *C. elegans* haplotype data from CeNDR. Swept strains that have at least one chromosome with equal or greater than 30% of swept haplotypes showed significantly higher lifetime fertility than divergent strains that have avoided the sweeps. We identified three QTL that underlie differences in lifetime fertility among the 121 *C. elegans* strains using single-marker based GWA mappings. Remarkably, across all three QTL, swept strains tend to have shared haplotypes and the reference alleles at peak markers. By contrast, divergent strains tend to have unswept haplotypes and the alternative alleles at peak markers. We also observed significant geographical differences in lifetime fertility between Hawaiian strains and strains from other parts of the world, likely because of the selective sweeps. We further mapped previous data using GWA mapping and linkage mapping and identified eight QTL underlying *C. elegans* fertility in different environments. Taken together, our results showed the diverse genetic basis of *C. elegans* fertility and suggest that higher fertility in most *C. elegans* strains could be caused by alleles that have recently swept throughout the world population.

### Genetically divergent strains have substantially lower fertility than swept strains

We measured lifetime fertility in 121 genetically distinct *C. elegans* strains. In our measurements (Figure 2A), the laboratory reference strain N2 (known as the Bristol strain) and a frequently used wild strain CB4856 (known as the Hawaii strain) had lifetime fertility of 308 and 237, respectively, with similar fertility values as reported previously (Hodgkin and Doniach 1997; Wegewitz *et al.* 2008; Andersen *et al.* 2014). The CB4856 strain had been considered the most genetically distant strain from the N2 strain for decades. In the last five years, researchers have collected and identified many genetically divergent *C. elegans* strains, some of which are more divergent from the N2 strain than the CB4856 strain is (Cook *et al.* 2017; Crombie *et al.* 2019; Lee *et al.* 2020). Most of these divergent strains were from Hawaii and showed none or rare evidence of the globally distributed swept haplotypes (Figure 1, Figure S1) (Crombie *et al.* 2019; Lee *et al.* 2020). In our fertility assays, we included many of these divergent strains. Under normal laboratory conditions, divergent strains showed significantly lower fertility than swept strains that have large blocks of swept haplotypes, suggesting that divergent strains have lower fitness than swept strains. The disadvantage in fertility of divergent strains was present from the beginning of the reproductive period throughout the peak. This lower fitness of divergent strains could have at least two possible explanations. First, laboratory conditions might favor swept strains over divergent strains. Standard laboratory conditions to culture *C. elegans* have been designed, modified, and improved based on the growth of the N2 strain (Brenner 1974), which is a swept strain. Most swept strains were from temperate zones (Andersen *et al.* 2012; Félix and Duveau 2012; Petersen *et al.* 2014; Richaud *et al.* 2018), such as Western Europe, whereas most divergent strains were isolated in the high elevation and cool temperature niches in the Hawaiian Islands (Crombie *et al.* 2019). The conditions of the natural habitats and the microenvironments in the niches of swept strains could be drastically different from niches of divergent strains. The closer the natural niche condition is to the laboratory condition, the higher fitness a swept strain might have. For instance, compared to N2, the strain CB4856 showed a clear thermal preference of approximately 17°C, which is lower than the canonical and the most typical *C. elegans* culture temperature of 20°C in the laboratory (Brenner 1974; Stiernagle 2006; Anderson *et al.* 2007). In a competition assay between two swept strains that isolated from locations with distinct climates, CX11314 (isolated at 20.9 °C) showed higher fitness than JU847 (isolated at 11.3°C) at both 15°C and 25°C, but JU847 grew better at 15°C than at 25°C (Evans *et al.* 2017). Divergent strains that were isolated from cool regions might exhibit higher fitness at temperatures lower than 20°C. The second explanation is that genetic variants at unknown loci directly caused differences in lifetime fertility between swept strains and divergent strains. The environment factors in our assays might have similar or minor influences on the fertility for both swept strains and divergent strains. The major differences in fertility between swept strains and divergent strains could be attributed to their genetic differences. For instance, because a *C. elegans* hermaphrodite produces 200 - 300 sperm in the late L4 stage before irreversibly switching to oogenesis to produce up to 1000 oocytes, the number of sperm limits fertility of self-fertilized hermaphrodites (Ward and Carrel 1979; Cutter 2004; Félix and Braendle 2010). Alleles at unknown loci in swept strains might lead to an increased number of sperm and thus a higher fertility than divergent strains. It is also possible that swept strains and divergent strains produce similar numbers of sperm, but divergent strains have higher embryonic lethality than swept strains. Because we quantified the viable offspring of each *C. elegans* strains as their fertility (See Methods), higher embryonic lethality could have caused the lower fertility in divergent strains. Although our GWA results might have mapped genomic regions underlying spermatogenesis or embryonic lethality, future efforts to quantify the numbers of sperm and fertilized embryos among wild *C. elegans* strains will help to further elucidate the differences in fertility among strains.

### Diverse QTL for lifetime fertility in different environments

We performed GWA mapping and identified three QTL on chromosome I, II, and V for lifetime fertility of *C. elegans,* which were grown on agar plates and fed *E. coli* OP50. The split of strains by genotypes at peak markers and the haplotypes of each strain in each QTL strongly suggest that the three QTL could be the genetic basis of different lifetime fertility between swept strains and divergent strains. The reference alleles and the most common haplotypes in each QTL, which provided the selective advantage of higher fertility, could have swept through the *C. elegans* population as these strains spread throughout the world. Under similar conditions, a previous study using linkage mapping and a large panel of RIAILs derived from the N2 and CB4856 strains have mapped fertility to QTL on chromosome II (2.6 Mb - 3.6 Mb) and X (4.6 Mb - 7.7 Mb) (Andersen *et al.* 2014). A laboratory-derived mutation in the gene *npr-1* from N2 was identified to have driven the QTL on chromosome X (McGrath *et al.* 2009; Andersen *et al.* 2014).

In liquid culture and fed the *E. coli* strain HB101, a new panel of *C. elegans* RIAILs with replacement of the N2 *npr-1* allele with the counterpart version from the CB4856 strain was used to map fertility to a QTL on chromosome IV (10.7 Mb - 12.8 Mb) by linkage mapping (Andersen *et al.* 2015). Using the same RIAILs panel but under three different liquid conditions (1% H2O, 1% DMSO, and 0.5% DMSO), we mapped fertility to seven QTL on chromosome II, IV, and V. In both DMSO conditions, the three QTL on chromosome IV recapitulated the QTL in the above study (Andersen *et al.* 2015); the two overlapping QTL on chromosome V overlapped with the QTL using our 121 wild strains grown in agar plates. We further used GWA to map previously published wild strain fertility data from liquid culture and 1% DMSO. A QTL linked to the selective sweeps located on the left arm of chromosome X was identified. Although *npr-1* is in the region of this QTL, the laboratory-derived N2 *npr-1* allele that is only found in the N2 strain could not drive this QTL because it is not found in wild strains.

As a complex life history trait, lifetime fertility could be influenced by many loci (Houle 1992). Under different conditions, GWA mappings identified QTL on chromosome I, II, V, and X; linkage mappings identified QTL on chromosome II, IV, V, and X. These results suggest that shared and separate loci in *C. elegans* genome control fertility in various environmental conditions. Because swept haplotypes shared among *C. elegans* strains might have driven all the QTL in GWA mappings, genetic variants in these swept haplotypes might be the beneficial alleles that swept through the *C. elegans* population.

### Potential adaptive alleles for *C. elegans* in temperate zones

The QTL for lifetime fertility using the 121 *C. elegans* strains also shared genomic regions with QTL on weather and climate variables related to natural habitats of 149 wild *C. elegans* strains (Evans *et al.* 2017). Two of the GWA mapping QTL for relative humidity were on chromosomes II and V, which overlapped with our QTL on chromosomes II and V, respectively. GWA mappings for three-year average temperature also located the same QTL just right of the center of chromosome V. We showed that *C. elegans* strains sampled from Europe and North America had similar lifetime fertilities, which were significantly larger than fertilities of Hawaiian *C. elegans* strains. Because Hawaii is in the tropical zone, *C. elegans* isolated from high elevation areas in Hawaii could have experienced high humidity and low temperatures in a much more stable climate in the long term than *C. elegans* in temperate zones. Alleles of swept strains in the shared QTL underlying lifetime fertility and climate variables could have enhanced the adaptability of *C. elegans* to variable humidity and temperatures in temperate zones along the *C. elegans* expansion out of the Pacific region (Andersen *et al.* 2012; Crombie *et al.* 2019; Lee *et al.* 2020). It is possible that, because of these adaptive alleles, the N2 strain showed no preference at these temperatures (Anderson *et al.* 2007).

Some Hawaiian strains, exclusively isolated at lower elevations closer to the coasts, exhibited admixture with non-Hawaiian populations, which might come from gene flow from outcrossing with immigrating swept strains from outside to Hawaii (Crombie *et al.* 2019). But compared to most non-Hawaiian strains, Hawaiian strains only contain, if any, small fractions of swept haplotypes. Of the 50 Hawaiian *C. elegans* strains used in this study, four strains are classified as swept strains, who have no more than two swept chromosomes (Figure S1). The alleles that increase lifetime fertility in swept strains might not contribute to higher fitness for *C. elegans* strains in Hawaii. In fluctuating environments in temperate zones, the randomly distributed and limited habitats might select for *C. elegans* that have higher fertility, although the high density of animals also facilitates dauer formation, which could underlie future survival success. Moreover, *C. elegans* populations in temperate zones also undergo bottlenecks in winter, from which dauers are more likely to survive. By contrast, Hawaiian *C. elegans* might not need to enter and stay in the dauer stage as often and long as non-Hawaiian *C. elegans* in temperate zones. Ample available habitats (*e.g*. rotting fruits) and the stable environment in Hawaii could lead to a higher survival rate for *C. elegans,* and thus lower fertility as a trade-off.

## ACKNOWLEDGMENTS

We would like to thank members of the Andersen Lab for helpful comments on the manuscript. G.Z. received support from the NSF-Simons Center for Quantitative Biology at Northwestern University (awards Simons Foundation/SFARI 597491-RWC and the National Science Foundation 1764421). J.D.M received support from a Northwestern Undergraduate Research Grant.

---

**Figure S1.**
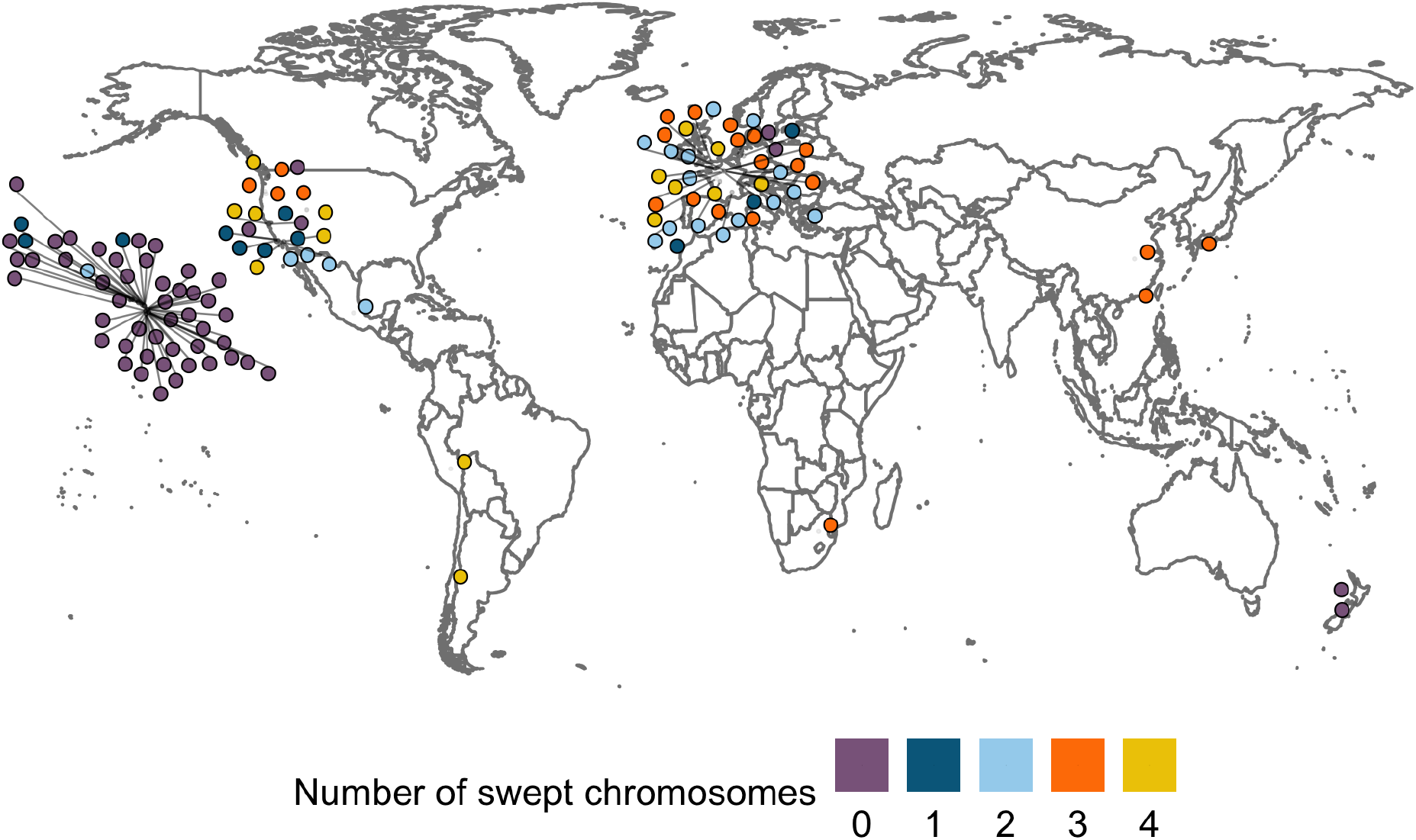
Global distribution of the 121 wild *C. elegans* strains used in this study. Each point corresponds to the isolation location and is colored by the number of swept chromosomes.

**Figure S2.**
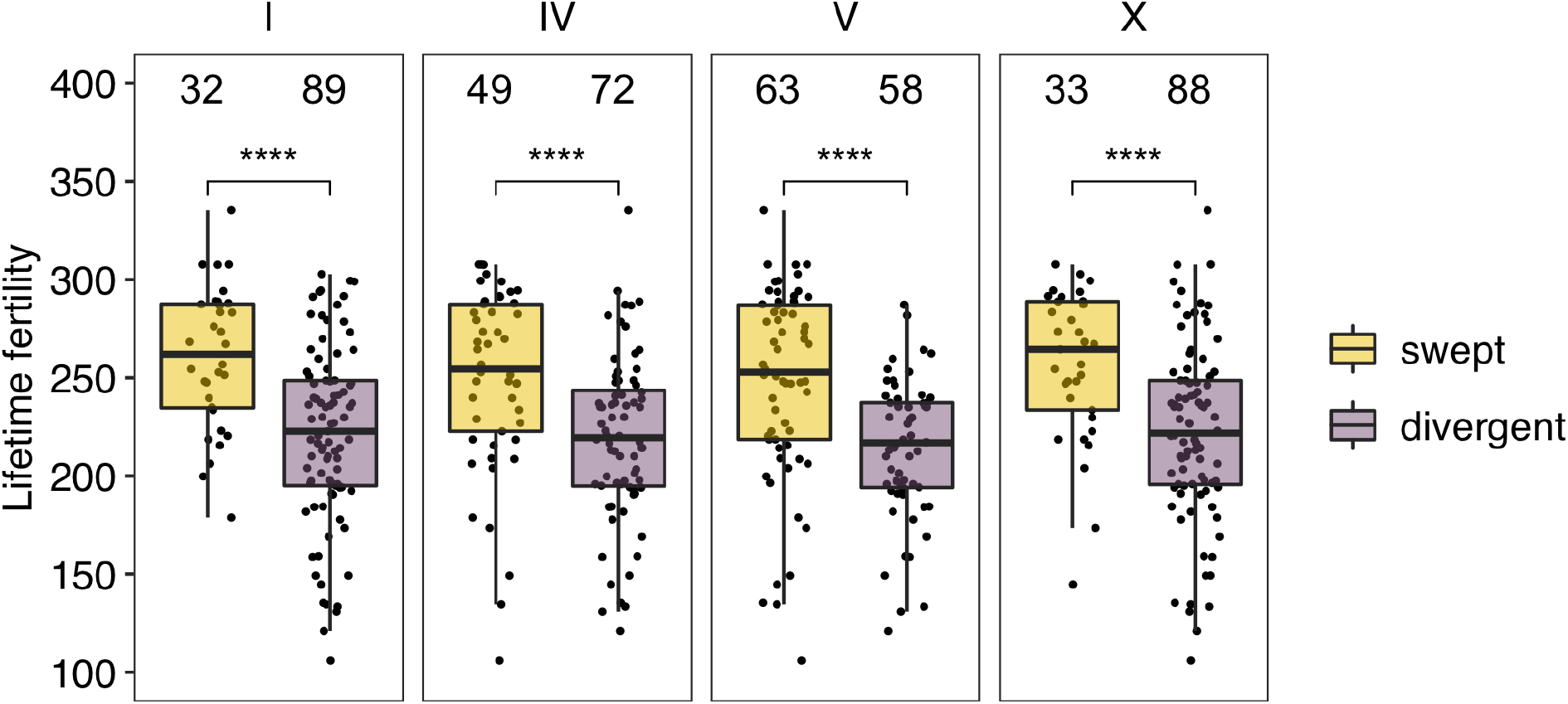
Comparisons of *C. elegans* lifetime fertility between swept groups (gold) and divergent groups (purple) classified by single chromosome are shown as Tukey box plots. Only swept chromosomes (I, IV, V, and X) are shown. For each chromosome (panel), strains with swept chromosomes were assigned to swept groups; strains with divergent chromosomes were assigned to divergent groups. Statistical significance was calculated using the Wilcoxon test, with *p*-values 3.7E-5, 5.5E-5, 6.5E-6, and 6.6E-5 in the comparisons by chromosomes I, IV, V, and X, respectively. Significance of each comparison is shown above each comparison pair (****: p-value ≤ 0.0001). The number of strains in each group is indicated above significance.

**Figure S3.**
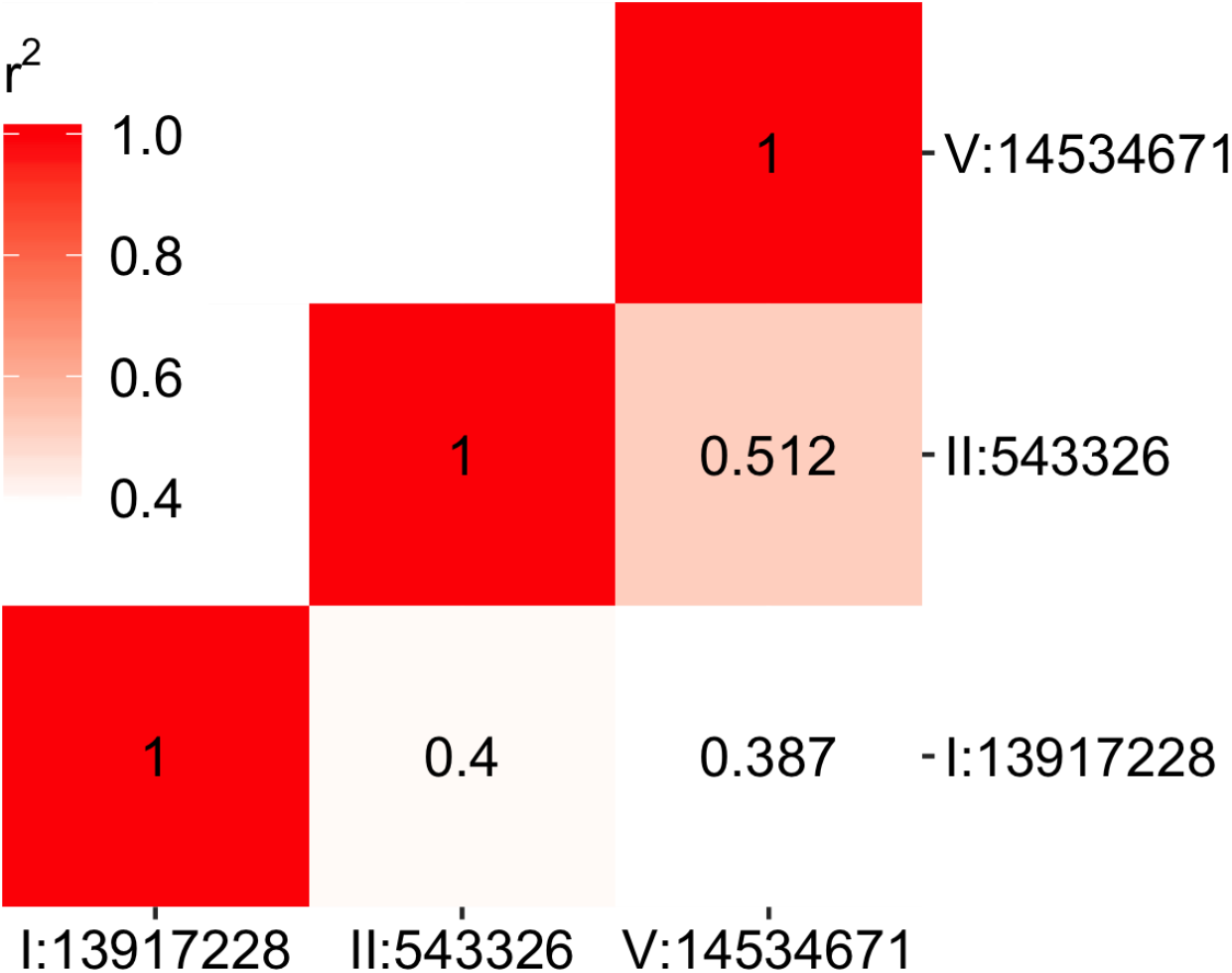
Linkage disequilibrium of QTL peak markers associated with *C. elegans* lifetime fertility is shown. Correlations (*r*^2^) between each marker pair are indicated in the tiles and are represented by the tile color.

**Figure S4.**
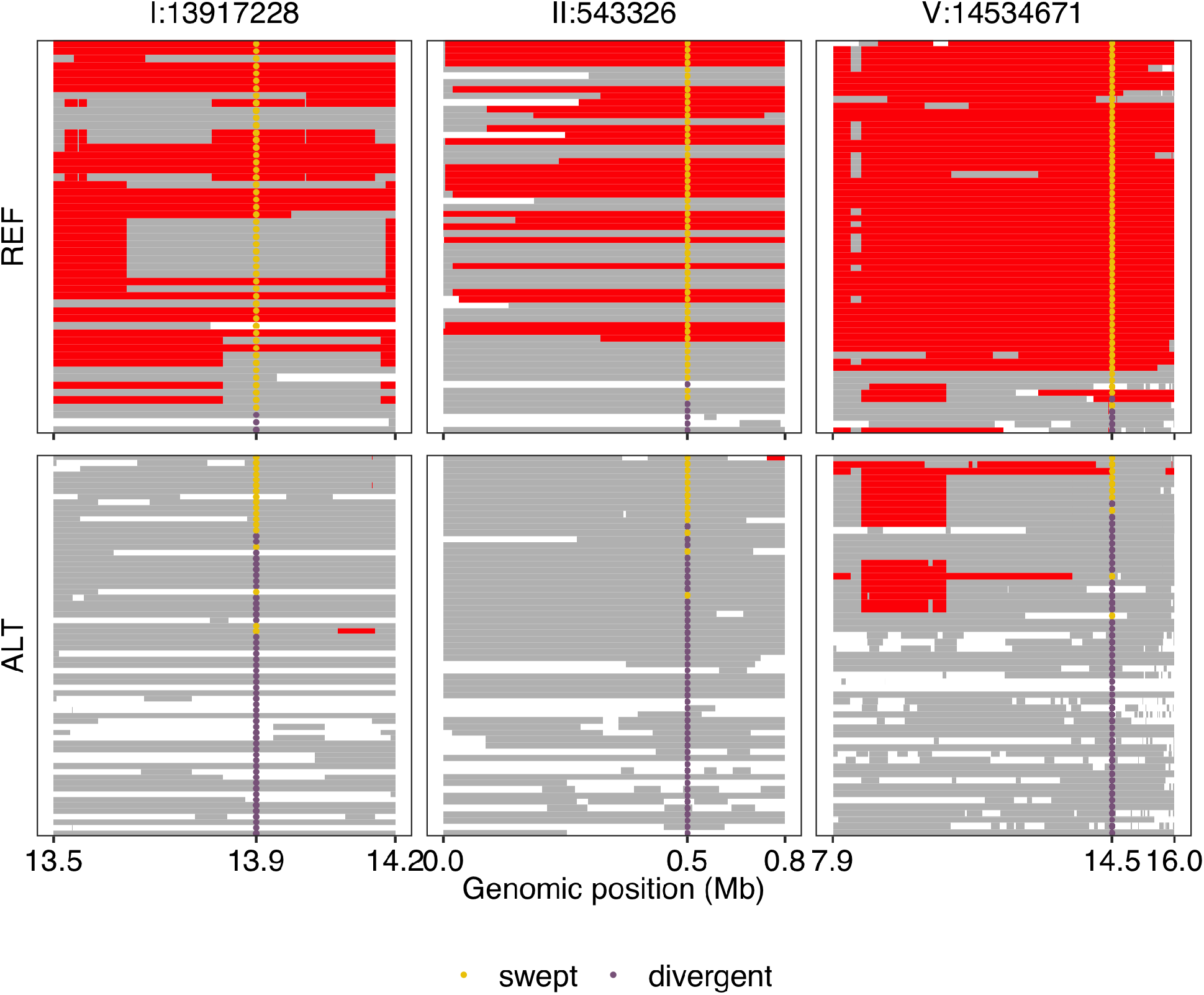
Sharing of haplotypes within QTL associated with lifetime fertility variation among 121 *C. elegans* strains is shown. Genomic regions of most common, rare and undetermined haplotypes are colored red, gray, and white, respectively. For each QTL (represented by peak markers on the top), strains were divided into REF (N2 reference alleles) panels or ALT (alternative alleles) panels by their genotypes at the peak markers as in Figure 3B. The genomic positions of each QTL are plotted on the x-axis. In the two panels of each QTL, each row on the y-axis represents one of the 121 strains and is ordered by their relative positions in Figure 1B. Swept strains and divergent strains are indicated as gold dots and purple dots, respectively, at the peak markers.

**Figure S5.**
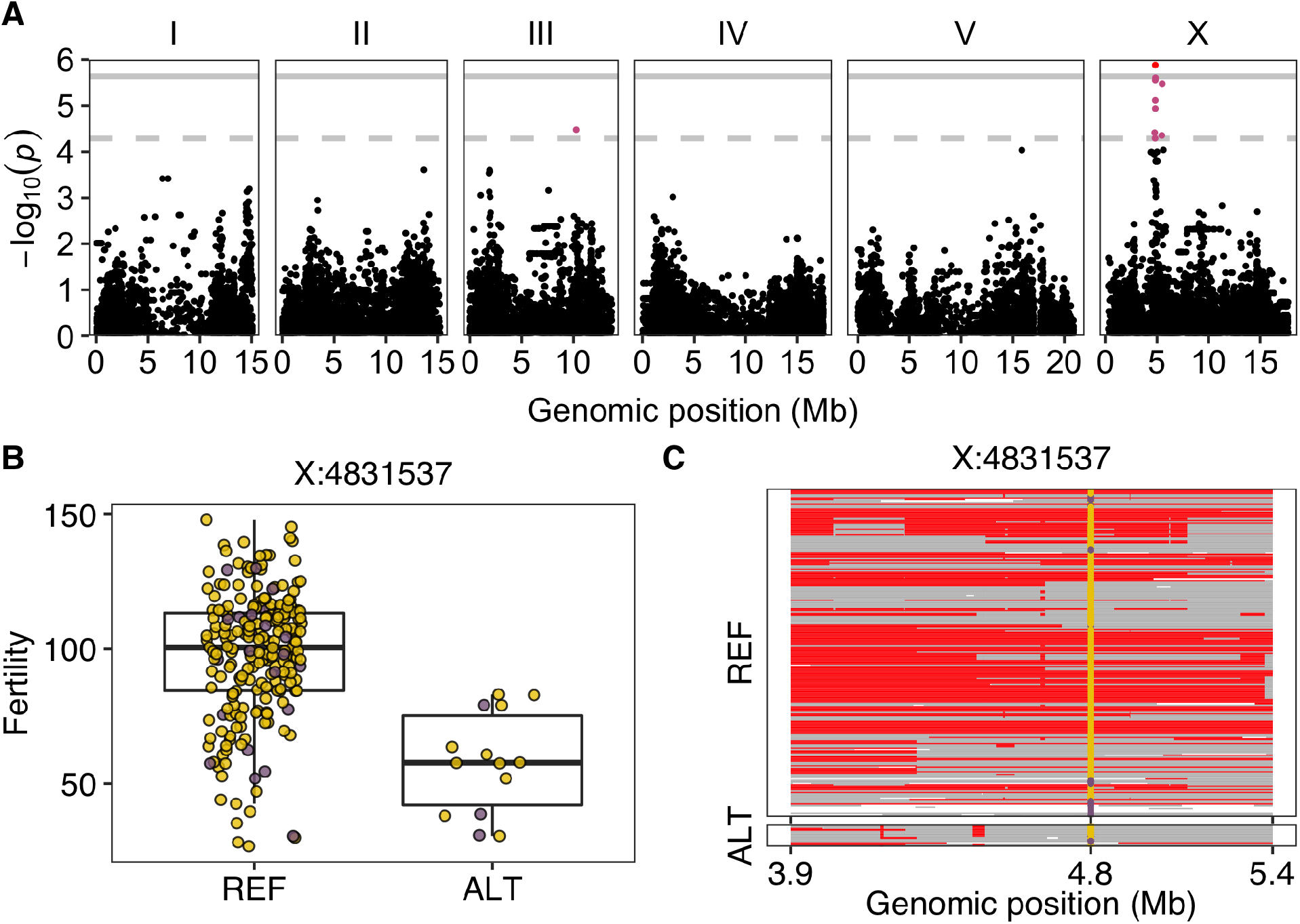
One QTL was identified in GWA mapping of *C. elegans* fertility variation in 236 strains. (A) Manhattan plot indicating GWA mapping results. Each point represents an SNV that is plotted with its genomic position (x-axis) against its −log_10_(p) value (y-axis) in mapping. SNVs that pass the genome-wide EIGEN threshold (the dotted gray horizontal line) and the genome-wide BF threshold (the solid gray horizontal line) are colored pink and red, respectively. (B) Tukey box plot showing fertility (norm.n) between strains with different genotypes at the peak marker position in the QTL. Each point corresponds to a *C. elegans* strain and is colored gold for swept strains and purple for divergent strains. On the x-axis, REF represents strains with the N2 reference allele and ALT represents strains with the alternative allele. (C) Sharing of haplotypes within the QTL is shown. Genomic regions of most common, rare and undetermined haplotypes are colored red, gray, and white, respectively. Strains were divided into REF (N2 reference alleles) panels or ALT (alternative alleles) panels by their genotypes at the peak markers as in (B). The genomic positions of the QTL are plotted on the x-axis. In the two panels of the QTL, each row on the y-axis represents one of the 236 strains, ordered as their relative positions in Figure 1B. Swept strains and divergent strains are indicated as gold dots and purple dots, respectively, at the peak markers.

**Figure S6.**
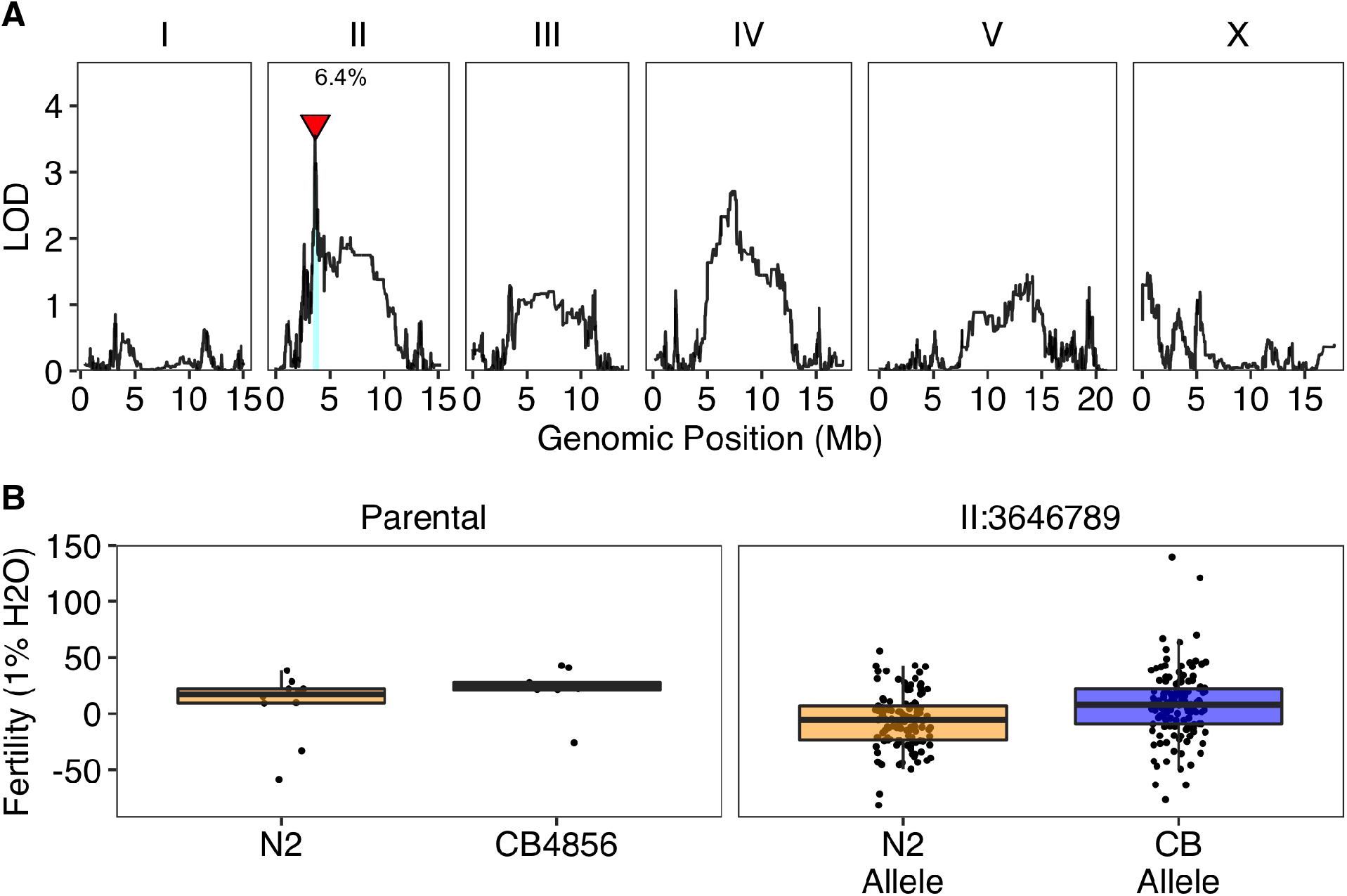
A QTL was identified using linkage mapping of *C. elegans* fertility (norm.n) in 1% water conditions. (A) Linkage mapping results of *C. elegans* fertility (norm.n) with RIAILs were shown with genomic position in Mb (x-axis) plotted against the logarithm of the odds (LOD) score (y-axis). The peak marker of the QTL on the left arm of chromosome II is indicated by a red triangle, next to which the percentage of the total phenotypic variance that can be explained by the QTL is shown. The 95% confidence interval of the QTL is shown by a blue rectangle. (B) Fertility (norm.n) is shown between the parents (N2 and CB4856) and between RIAILs split by genotype at the peak marker of the QTL. Each dot in the parental panel represents one of the replicates. Each dot in the QTL panel corresponds to a unique recombinant strain. Strains with the N2 allele are colored orange, and strains with the CB4856 allele are colored blue.

**Figure S7.**
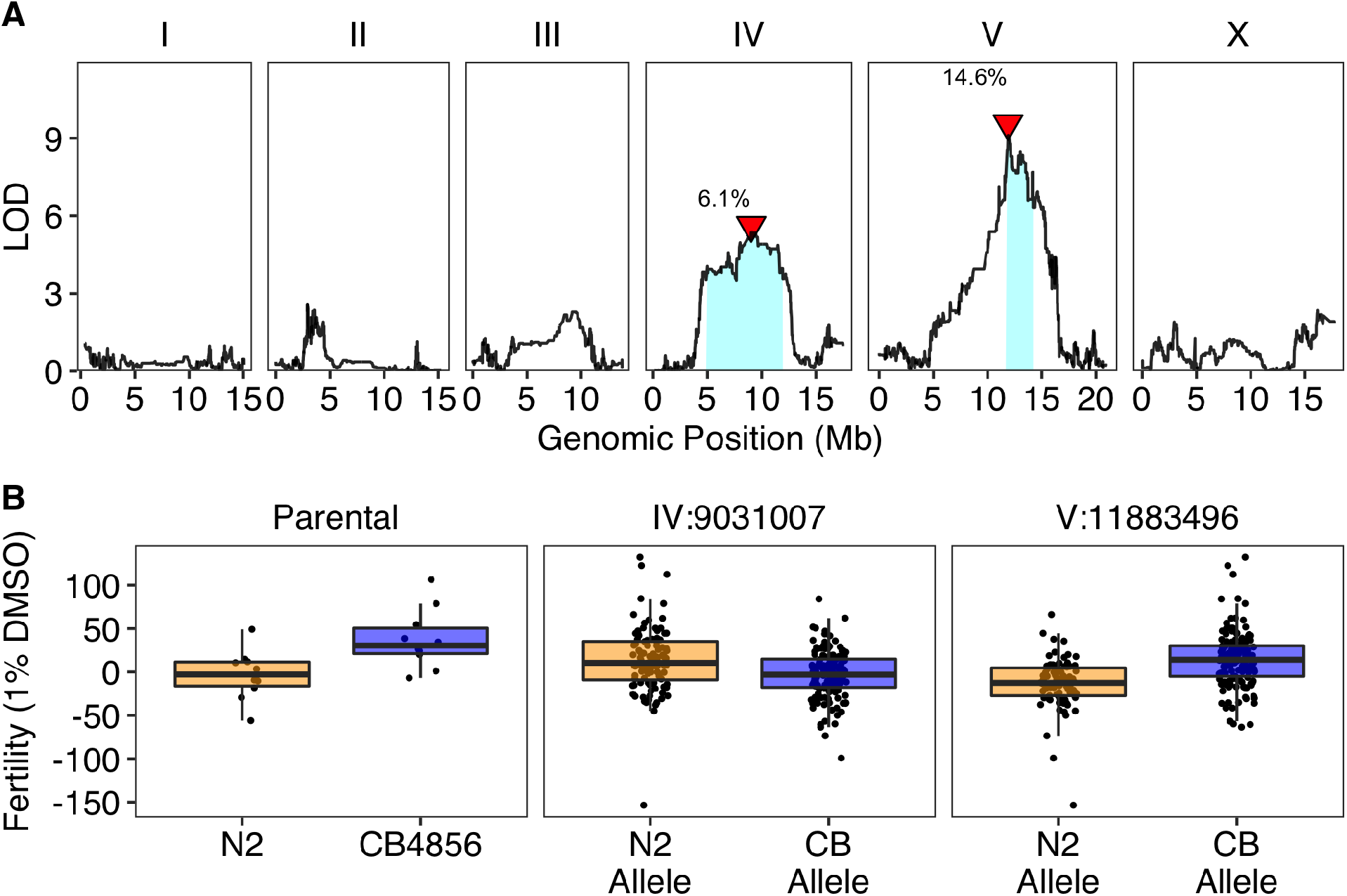
Two QTL were identified using linkage mapping of *C. elegans* fertility (norm.n) in 1% DMSO conditions. (A) Linkage mapping results of *C. elegans* fertility (norm.n) with RIAILs were shown with genomic position in Mb (x-axis) plotted against the logarithm of the odds (LOD) score (y-axis). The peak markers of QTL are indicated by red triangles, next to which the percentages of the total phenotypic variance that can be explained by the QTL are shown. The 95% confidence interval of each QTL is shown by a blue rectangle. (B) Fertility (norm.n) is shown between the parents (N2 and CB4856), and between RIAILs split by genotype at the peak marker for each QTL. Each dot in the parental panel represents one of the replicates. Each dot in each of the QTL panels corresponds to a unique recombinant strain. Strains with the N2 allele are colored orange and strains with the CB4856 allele are colored blue.

**Figure S8.**
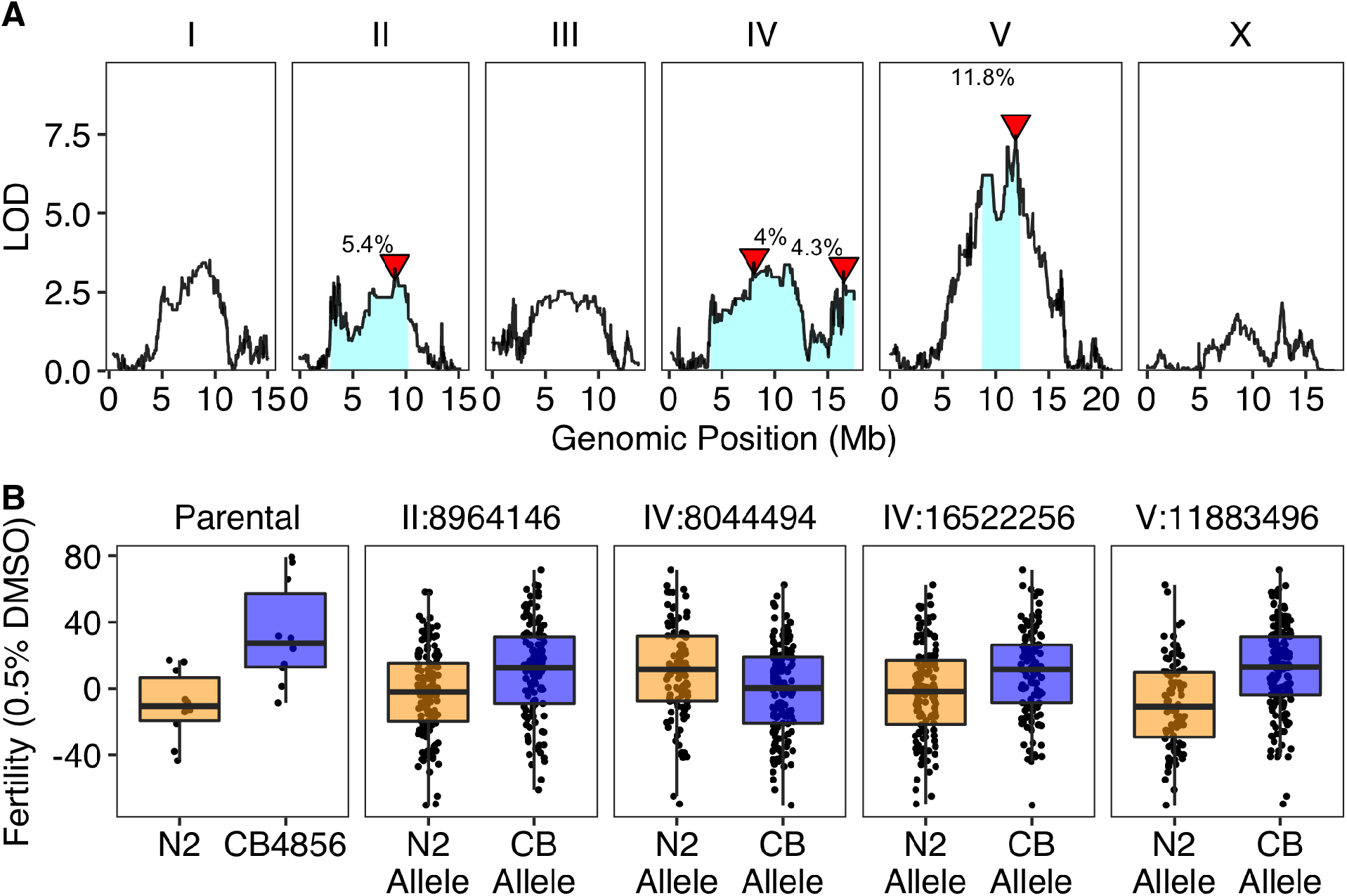
Four QTL were identified using linkage mapping of *C. elegans* fertility (norm.n) in 0.5% DMSO conditions. (A) Linkage mapping results of *C. elegans* fertility (norm.n) with RIAILs were shown with genomic position in Mb (x-axis) plotted against the logarithm of the odds (LOD) score (y-axis). The peak markers of QTL are indicated by red triangles, next to which the percentages of the total phenotypic variance that can be explained by the QTL are shown. The 95% confidence interval of each QTL is shown by a blue rectangle. (B) Fertility (norm.n) is shown between the parents (N2 and CB4856), and between RIAILs split by genotype at the peak marker for each QTL. Each dot in the parental panel represents one of the replicates. Each dot in each of the QTL panels corresponds to a unique recombinant strain. Strains with the N2 allele are colored orange, and strains with the CB4856 allele are colored blue.

